# MET-2, a SETDB1 family methyltransferase, coordinates embryo events through distinct histone H3 methylation states

**DOI:** 10.1101/429902

**Authors:** Beste Mutlu, Huei-Mei Chen, David H. Hall, Susan E. Mango

## Abstract

During the first hours of embryogenesis, formation of higher-order heterochromatin coincides with the loss of developmental potential. Here we examine the relationship between these two processes, and we probe the determinants that contribute to their onset. Mutations that disrupt histone H3 lysine 9 (H3K9) methyltransferases reveal that the methyltransferase MET-2 helps terminate developmental plasticity, likely through mono- and di- methylation of H3K9 (me1/me2), and promotes heterochromatin formation, likely through H3K9me3. We examine how MET-2 is regulated and find that methylated H3K9 appears gradually and depends on the accumulated time of embryogenesis. H3K9me is independent of zygotic genome activation or cell counting. These data reveal how central events are synchronized during embryogenesis and distinguish distinct roles for different H3K9 methylation states.

**Summary Statement:** During early embryogenesis, heterochromatin formation and loss of developmental plasticity are coordinately regulated by distinct Histone H3 Lysine 9 (H3K9) methylation states, by the methyltransferase MET-2.

## Introduction

Upon fertilization, embryos initiate developmental events to control cell division, pluripotency, differentiation, reorganization of the nucleus, onset of transcription and repositioning of cells into germ layers that ultimately generate organs and tissues. These processes are coordinated in time and space. In *C. elegans*, for example, the reorganization of cells during gastrulation coincides with loss of developmental plasticity, the formation of heterochromatin and a surge of transcription. While the timing of these events has been defined (Mango, 2009; Mutlu et al., 2018; Robertson et al., 2004; Seydoux and Fire, 1994; Sulston et al., 1983; Yuzyuk et al., 2009), little is known about the processes that regulate the timing or synchronization of different events (Detwiler et al., 2001; Mutlu et al., 2018; Yuzyuk et al., 2009). To begin to address this issue, we have examined whether early embryonic events are contingent upon each other or regulated independently.

*C. elegans* embryos, like other those of other animals, are born pluripotent and lack higher-order heterochromatin, the portion of the genome that carries repetitive DNA and is less transcriptionally active (Mutlu et al., 2018). As embryos transition towards gastrulation, they lose developmental potential and generate heterochromatin (Mutlu et al., 2018). In cultured mammalian cells, the enzymes that promote heterochromatin formation also inhibit reprogramming into embryonic stem cells, suggesting a possible link between these processes (Gaspar-Maia et al., 2009; Kang, 2014; Soufi et al., 2012; Zaret and Mango, 2016). For example, in mammals, the H3K9 methyltransferase SETDB1 both promotes heterochromatin formation and dampens reprogramming efficiency (Becker et al., 2015). However, SETDB1 is responsible for multiple H3K9 modifications, making it difficult to determine whether different methylated states have distinct roles. Conversely, in many mammals, multiple enzymes contribute to a given modification, which complicates studying the function of a single histone modification. We use *C. elegans*, which enabled us to dissect the roles of different H3K9 methylated states. *C. elegans* MET-2 is homologous to SETDB1 (Andersen and Horvitz, 2007; Poulin et al., 2005) and is responsible for virtually all embryonic H3K9me1/me2, but has only partial effects on H3K9me3; SET-25 is necessary for virtually all H3K9me3, but not H3K9me1/me2 (Towbin et al., 2012). The separation of enzymatic activities provides an opportunity to determine which modifications govern which processes. Studying an intact animal also enables us to establish the temporal order of events in relation to developmental milestones.

The formation of heterochromatin during *C.elegans* embryogenesis depends on MET-2 (Mutlu et al., 2018). Early embryos have almost no H3K9me2 and low levels of H3K9me1 and H3K9me3. By transmission electron microscopy, they lack heterochromatin. As embryos develop, MET-2 becomes enriched in the nucleus, promotes an increase in all three methylated forms of histone H3 and generates heterochromatin (Mutlu et al.,2018). It is unknown if MET-2 also regulates developmental potential, nor are the separate roles of mono, di and tri H3K9me understood. Here we examine these different roles by studying MET-2 and SET-25, and we investigate how MET-2 and H3K9 methylation are regulated in early embryos.

## Results

### MET-2 promotes loss of plasticity

We took advantage of the Cell Fate Challenge assay (CFC) (Horner et al., 1998; Mango, 2009) to test the importance of *met-2* for developmental plasticity (**Figure 1**). *C. elegans* embryos develop within 14 hours, with initiation of zygotic transcription at the 4-cell stage and gastrulation at the 28-cell stage (Schauer and Wood, 1990; Seydoux and Fire, 1994; Sulston et al., 1983). Prior to gastrulation, cells are developmentally plastic, and their normal pattern of development can be reprogrammed by ectopic expression of selector genes, but this flexibility is lost during gastrulation (Djabrayan et al., 2012; Fukushige and Krause, 2005; Gilleard and McGhee, 2001; Horner et al., 1998; Mango, 2009; Priess and Thomson, 1987; Sulston et al., 1983; Wood, 1991; Zhu et al., 1998).

Embryos were induced to alter their cell identity and acquire muscle fate by ectopic expression of *hlh-1/MyoD* under control of the heat-shock promoter (*HS::hlh-1*; (Fukushige and Krause, 2005)). We focused on the 80–100-cell stage (mid-gastrulation), when wild-type embryos have an intermediate response to the CFC assay, thereby providing a sensitive assay (**Figure 1A-B**). In addition, H3K9me2 is readily visible at this developmental stage in wild-type embryos but absent from *met-2* mutants (Mutlu et al., 2018). The foregut marker PHA-4 was used to identify cells that retained their endogenous identity and resisted exogenous HLH-1, and muscle paramyosin was used to track conversion to muscle fate (Horner et al., 1998; Yuzyuk et al., 2009). We concentrated on endogenous fate markers because exogenous reporters can sometimes lead to spurious expression or regulation (Jiao et al., 2018; Mango, 2007). As a control, we examined *hlh-1* mRNA expression before and after heat shock and observed no difference in induction between wild-type and mutant embryos (**Figure 1D**).

When challenged with *HS::hlh-1*, 35% of 100-cell-stage wild-type embryos converted to muscle fate completely, consistent with previous studies (Fukushige and Krause, 2005; Mango, 2009; Yuzyuk et al., 2009). In *met-2* mutants, 54% of embryos responded to *HS::hlh-1* with a complete cell-fate transformation (**Figure 1C**, 100-cell stage, n=4, > 100 embryos each, p=0.008). This result suggests that, normally, *met-2* promotes loss of plasticity during early gastrulation. By the ∼200 cell stage, none of the wild-type or *met-2* mutant embryos remained fully plastic (**Figure 1C**). Moreover, both wild-type and *met-2* mutant embryos had a similar number of cells that resisted changing fate, indicating that *met-2* mutants eventually terminate plasticity (**Supplementary Figure 1**). The data reveal that *met-2* restricts developmental plasticity during gastrulation but is not absolutely required. The partial requirement for *met-2* may reflect that other regulators also control developmental plasticity in the embryo (Djabrayan et al., 2012; Joshi et al., 2010; Yuzyuk et al., 2009).

H3K9me1/me2 exist in the embryo and, in addition, are used to produce H3K9me3 by SET-25 (Towbin et al., 2012). We wondered whether either the plasticity or the heterochromatin phenotype of *met-2* could be explained by its role in generating H3K9me3. First, we examined developmental plasticity with the CFC assay. Inactivation of *set-25* lead to the opposite result as *met-2*: only 20% of *set-25* embryos were developmentally plastic, about half of wild-type (**Figure 1C**, n=3 experiments, > 60 embryos each, p=0.011). This result demonstrated that H3K9me3 was not required to terminate plasticity during gastrulation. Rather, developmental plasticity correlated with the level of H3K9me2, which is low in *met-2* mutants and high in *set-25* mutants relative to wild-type **(Supplementary Figure 2)**.

To test the importance of H3K9me2, we examined *met-2; set-25* double mutants, which lack mono, di and tri H3K9me (Garrigues et al., 2014; Towbin et al., 2012). The double mutants had prolonged plasticity (**Figure 1C**, n=2, 30 embryos each, p=0.045), revealing that the reduced plasticity of *set-25* mutants required wild-type *met-2* activity. A simple hypothesis to explain these results is that H3K9me1 and/or H3K9me2 is required to terminate developmental plasticity but H3K9me3 is not. To gain insight into genes regulated by H3K9me1 and H2K9me2, we examined modENCODE data for genes bearing either of these marks. Among genes with peaks of H3K9me1, there was enrichment for GO terms “*intracellular signaling cascade*,” and regulation of “*developmental process*” and “*growth*” (Figure 1E). For H3K9me2, there was enrichment for “*cell-fate determination*” and “*embryonic pattern specification*,” which included genes expressed in the pre-gastrula embryo (e.g. *par-1, par-4, mbk-2* and others). These GO terms were not enriched for H3K9me3, possibly explaining the distinct phenotypes of *met-2* vs. *set-25*.

**Figure 1.**
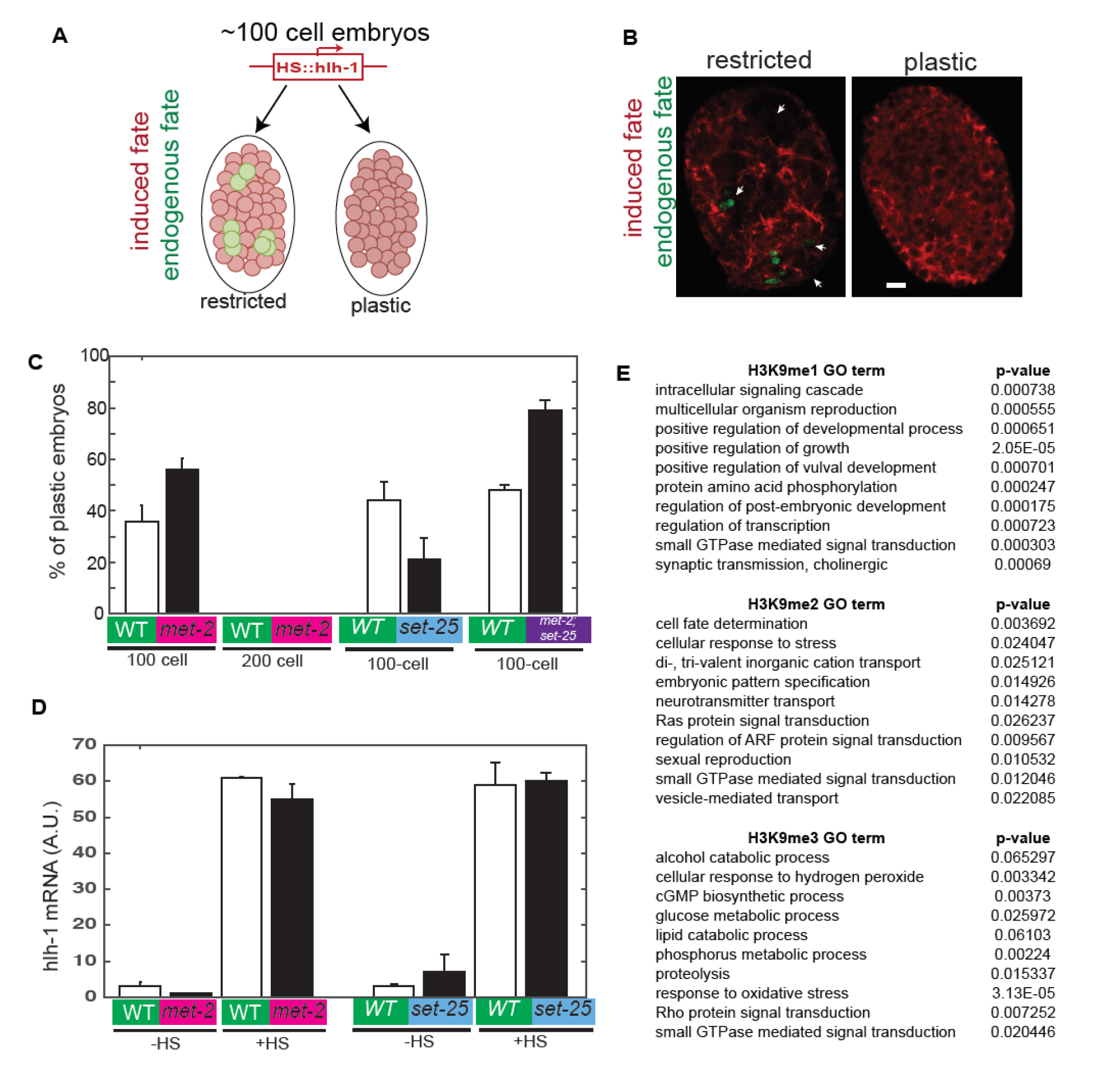
MET-2 promotes loss of developmental plasticity at mid-gastrulation. A. Cell Fate Challenge Assay (CFCA). Muscle fate regulator *hlh-1/MyoD* is induced in somatic cells. Terminally differentiated embryos stained for induced or endogenous fate markers (muscle paramyosin, red, or foregut PHA-4, green, respectively (Fukushige and Krause, 2005; Horner et al., 1998)). B. Terminally differentiated embryos. Arrowheads depict cells lacking paramyosin, some of which express PHA-4. Scale bar, 5μm. C. Percentage of *wild-type* (n=109) vs. *met-2* (n=136) or *wild-type* (n=64) vs. *set-25* (84) *or wild-type* (n=31) *vs. met-2; set-25* (n=38) embryos scored as developmentally plastic based on absent PHA-4 and ubiquitous paramyosin. Note that MET-2 promotes loss of plasticity, while SET-25 prevents it. D. *hlh-1/MyoD* mRNA levels normalized to *eft-3* in *wild-type* or mutant embryos, with (+HS) or without (-HS) heat-shock induction. CFCA results are not due to differential induction of hlh-1 in wild-type vs. mutants. E. GO Term Analysis for genes enriched for H3K9 methylation. Genes associated with H3K9me2 are involved in regulating plasticity, whereas H3K9me1 and H3K9me3 are not.

### H3K9me3 methyltransferase SET-25 is required for heterochromatin formation

Next, we examined the role of MET-2 and SET-25 for higher-order heterochromatin, as visualized by transmission electron microscopy (TEM). Wild-type nuclei from young embryos were predominantly electron translucent whereas nuclei from older embryos contained many electron dense regions (EDRs) (**Figure 2A**), as observed previously (Mutlu et al., 2018). EDRs represent heterochromatin regions based on morphology and on the enrichment for silencing marks such as H3K9me3 (Rübe et al., 2011). In *met-2* mutants, EDRs are paler than normal and delayed in their appearance (Mutlu et al., 2018). In *set-25* mutants, we did not detect any EDRs through the 200-cell stage, a phenotype that was stronger than *met-2* mutants (**Figure 2A**). As a control for TEM fixation and imaging, we examined the cytosol of wild-type and *set-25* mutants, using a threshold value to define electron dense regions. Both genotypes had darkly stained cytosolic regions (**Figure 2B**). These data show that disruption of EDR formation tracks with H3K9me3: intermediate EDR formation in *met-2* mutants correlates with intermediate H3K9me3 levels and absent EDRs in *set-25* mutants with absent H3K9me3. These results demonstrate that neither H3K9me3 nor the formation of large-scale, higher-order heterochromatin is required to terminate developmental plasticity. The data suggest that coordination between the termination of plasticity and generation of heterochromatin relies on distinct readouts from MET-2: formation of heterochromatin from H3K9me3 and termination of plasticity by H3K9me1/me2 (**Figure 2C**).

**Figure 2.**
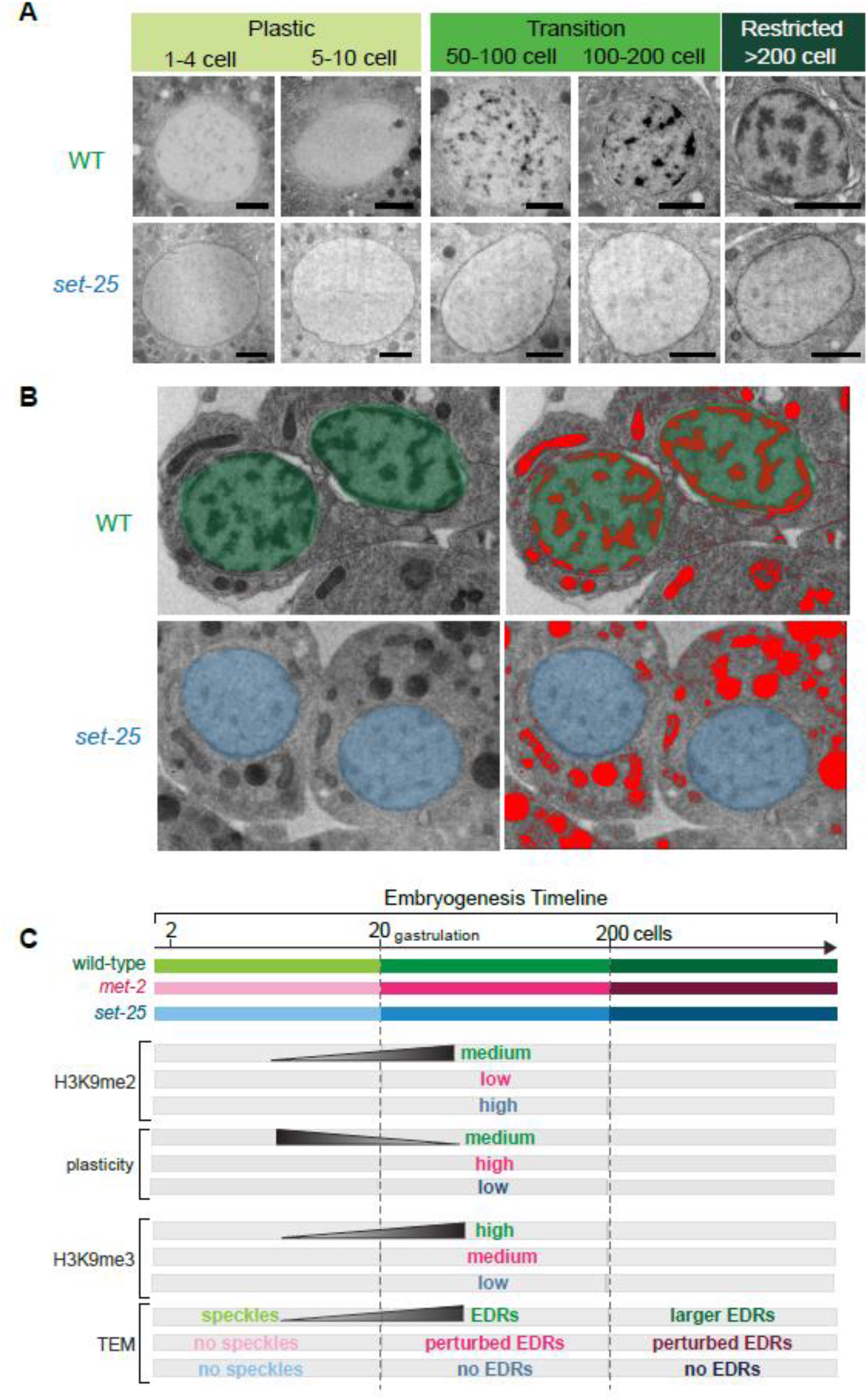
H3K9me3 methyltransferase SET-25 is required for heterochromatin formation. A. TEM of representative nuclei from WT or *set-25* mutant embryos. Scale bars 1μm. For *set-25* embryos: N=2, 14 “Plastic”, 10 “Transition” and 5 “Restricted” embryos, with > 20 nuclei for each group. B. Contrast of the cytosol in WT or *set-25* mutants and intensity thresholding (red) to define electron-dense structures. C. Summary of results in WT, *met-2* and *set-25* mutant embryos. Embryonic stages analyzed in this study are early (<20 cell), mid (21–200 cell) and late (>200 cell).

### H3K9 de-methylation is not a timer for establishing H3K9me2 domains at gastrulation

The above data suggest that MET-2 helps coordinate two critical events in the early embryo: loss of developmental plasticity by H3K9me1/me2 and generation of higher-order heterochromatin by H3K9me3. Both of these processes occur during gastrulation as H3K9me accumulates, raising the question of what mechanisms regulate H3K9me temporally (**Figure 3A**). Since MET-2 acts upstream of SET-25 (Towbin et al., 2012) and H3K9me2 increased most dramatically from fertilization to gastrulation (Mutlu et al., 2018), we focused on the timing of H3K9me2 onset.

Initially, we wondered if an H3K9me2 demethylase might act in early embryos to remove H3K9me2. To test this idea, we examined a mutant for the H3K9me2 demethylase *jmjd-1.2* (Kleine-Kohlbrecher et al., 2010). In *jmjd-1.2(tm3713)* mutants that don’t produce JMJD-1.2 (Myers et al., 2018), H3K9me2 was extremely low in early embryos, like wild-type, and was established normally at gastrulation (**Figure 3B-D**). These data suggested that histone de-methylation is not part the H3K9me2 timer and led us to focus on MET-2.

**Figure 3.**
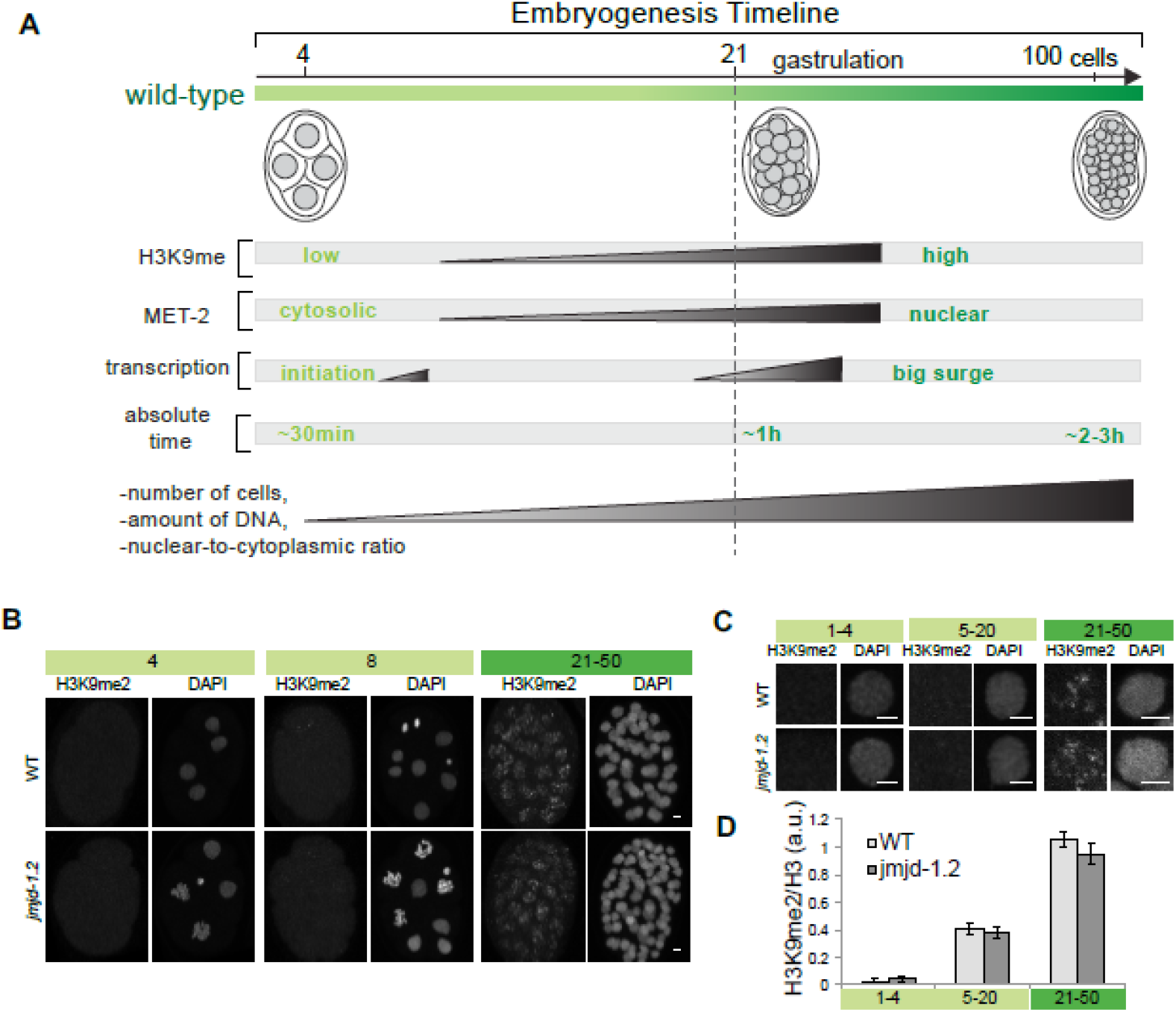
H3K9me1/me2 methyltransferase MET-2 is regulated during embryogenesis to form heterochromatin. A. Diagram showing a timeline for *C. elegans* embryogenesis and a summary of changes that accompany differentiation. With each cell division and passing time, the total number of cells, amount of DNA per embryo and the nuclear-to-cytoplasmic ratio per cell increases. The transition from light to dark green represents initiation of gastrulation around the 28-cell stage. Zygotic transcription is initiated at the 4-cell stage, but a big wave coincides with gastrulation (Levin et al., 2012; Schauer and Wood, 1990; Storfer-Glazer and Wood, 1994). All of these changes are candidates for dictating the timing of H3K9me2. C. H3K9me2 levels in wild-type vs. H3K9me2 demethylase *jmjd-1.2* mutants at pregastrula (4-cell and 8-cell stage) and gastrula (21–50 cell) stages. D-E. Single nuclei in interphase showing H3K9me2 levels in wild-type vs. *jmjd-1.2* mutants. Quantitation of H3K9me2 levels per nucleus. Early *jmjd-1.2* embryos do not have increased H3K9me2.

### Zygotic transcription is not rate-limiting for H3K9 di-methylation

The onset of gastrulation and surge in H3K9me2 is accompanied by a big wave in zygotic transcription (Baugh, 2003; Edgar et al., 1994; Hsu et al., 2015; Levin et al., 2012; Storfer-Glazer and Wood, 1994; Yuzyuk et al., 2009). We hypothesized that *met-2* or its cofactors could be activated in the embryo as zygotic transcription begins (**Figure 4A**, Model 1). In addition, transcription elongation could be rate-limiting for recruiting MET-2 to specific loci through interactions with the RNAi machinery ((Guang et al., 2010), **Figure 4A**, Model 2). We tested these models in two ways. First, we blocked zygotic transcription and examined whether the onset of H3K9me2 was disturbed. Second, we tested whether a zygotic copy of *met-2* could rescue H3K9me2 in the absence of maternal MET-2. For these experiments, we were able to compare H3K9me2 levels across different genotypes by including wild-type embryos marked with a fluorescent tag as an on-slide control. Control embryos and test embryos were processed together on the same slide and imaged with identical settings (**Figure 4B**).

To block transcription, we inactivated the canonical TFIID subunit, *taf-6.2*, with a temperature-sensitive mutation *(ax514)* (Bowman et al., 2011), and a core subunit of RNA Polymerase II, *ama-1*, by RNAi. To assess the strength of the block on transcription, we stained embryos with the H5 Polymerase II antibody against the phosphorylated Ser2 on the CTD domain, which is a hallmark of transcriptional elongation (Bregman et al., 1995; Seydoux and Dunn, 1997). Under our conditions, H5 Polymerase II signal was undetectable after blocking zygotic transcription, whereas non-specific staining of P-granules by the H5 antibody served as a positive control. (**Figure 4C, D**). Despite the lack of detectable H5, H3K9me2 levels were identical to wild-type embryos (**Figure 4C, E**), suggesting that neither zygotic genes nor elongating Polymerase II were rate-limiting for H3K9me2 onset.

The transcriptional block suggested that MET-2 and its regulators must be contributed maternally and independent of zygotic transcription. To test this idea further, we examined RNA expression of *met-2* and its binding partners *lin-65* and *arle-14* (Mutlu et al., 2018), which are required for H3K9me2. *met-2, lin-65* and *arle-14* transcripts were abundant in early embryos, before the initiation of zygotic transcription (Levin et al., 2012; Seydoux and Fire, 1994; Tintori et al., 2016), revealing that these RNAs were deposited by the mother. To test if zygotic MET-2 could rescue H3K9me2 in embryos, we mated *met-2* mutant mothers with wild-type males and analyzed the progeny, which carried a single zygotic copy of the wild-type *met-2* locus. Zygotic *met-2* could not restore H3K9me2 to mutant embryos (**Figure 4F**). Similarly, neither zygotic *lin-65* nor zygotic *arle-14* could rescue H3K9me2 from maternally mutant mothers (Figure 3.2G). Moreover, when we mated wild-type mothers with *met-2::gfp* males, the resulting *met-2::gfp* progeny never expressed GFP (**Figure 4H,I**). These results demonstrate that maternally deposited MET-2 and its binding partners are responsible for establishing H3K9me2 in embryos, and that the onset of transcription cannot account for the timing of H3K9me2 deposition.

**Figure 4.**
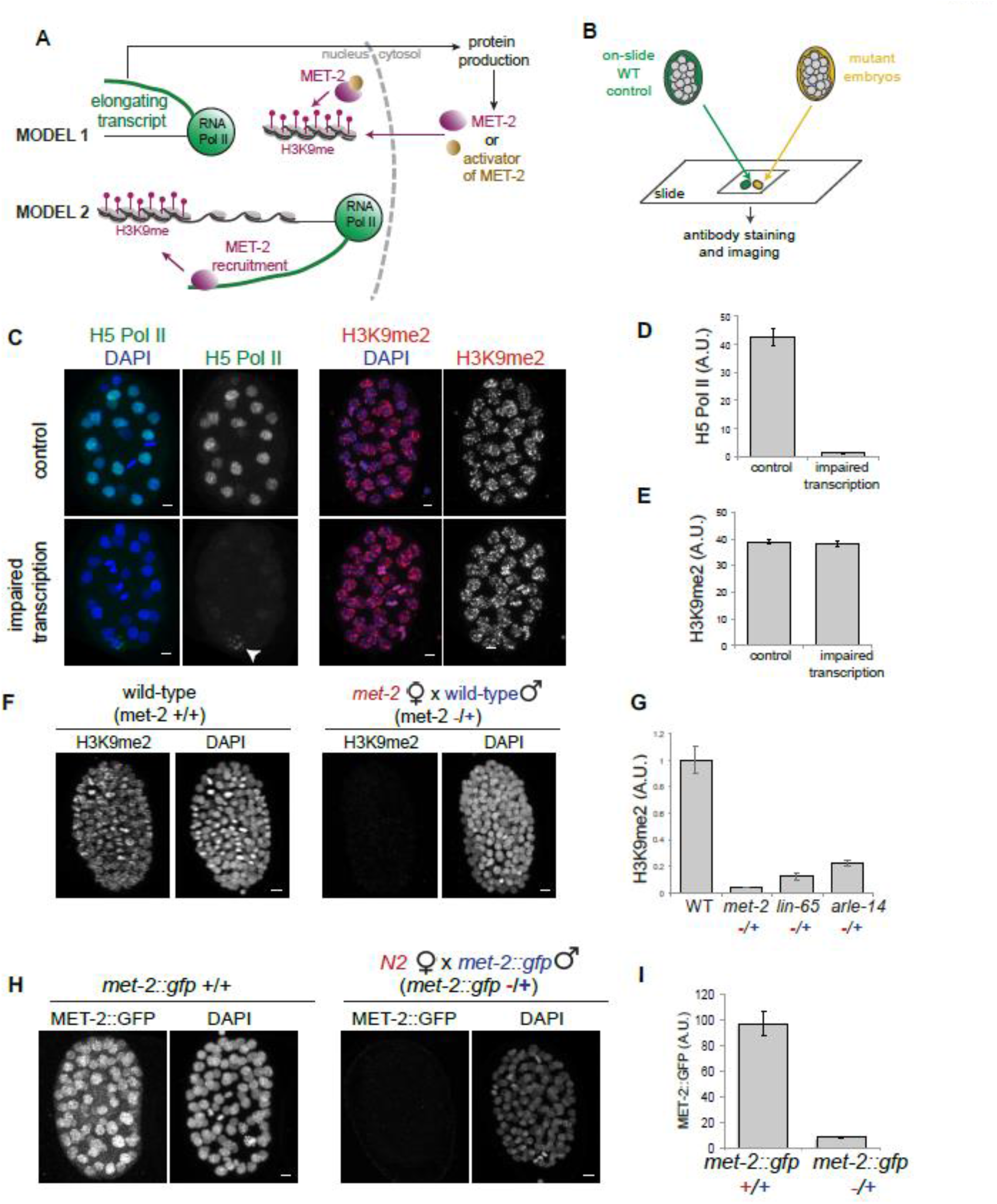
Zygotic transcription is not rate-limiting for H3K9 di-methylation. A. Two models describe how initiation of zygotic transcription might be important for dictating the timing of H3K9me2. Model 1: Transcription could be important for the production of MET-2 protein or an activator of MET-2. Model 2: Transcription itself might be rate-limiting to recruit MET-2 to chromatin (Bühler and Moazed, 2007; Guang et al., 2010) B. Experimental design. Wild-type embryos (green) containing a single copy ZEN-4::GFP or HIS-72::mCHERRY marker were analyzed on the same slide as unlabeled mutant embryos (yellow). This approach controlled for staining variability, thereby enabling an accurate comparison between different genotypes. C. Gastrula embryos stained with an antibody that detects the elongating form of RNA Polymerase II (H5; green) or H3K9me2 (red). Scale bar 2 μm. Transcription was blocked by combining *ama-1* RNAi with the *ax514* mutation in the initiation factor *taf-6.2*. White arrow points to non-specific P-granule (Wang and Seydoux, 2014) staining by the H5 antibody and serves as a staining control. D-E. Quantitation of mean signal intensity per nucleus for Pol II and H3K9me2 in wild-type vs. transcriptionally impaired embryos on the same slide. Note that transcription elongation is undetectable but H3K9me2 levels are unaffected. F-G. H3K9me2 levels in gastrula embryos that lack maternal *met-2, lin-65* or *arle-14* but contain a paternal copy. The genotype of the mother is highlighted in red, the father’s in blue. Zygotic copies of these genes cannot rescue H3K9me2 in the embryo. H-I. MET-2::GFP levels in gastrula embryos that lack a maternal copy of the construct, but inherit a paternal copy. Note that the paternal copy of MET-2 is not expressed in the embryo and maternal MET-2 sufficient for all H3K9me2 during embryogenesis.

### Length of interphase, but not cell counting mechanisms, dictates timing of H3K9me2

The transcription block ruled out some expected mechanisms for H3K9me2 onset. To gain a broader perspective on regulation of H3K9me onset, we considered alternative models. Embryonic timing has been studied extensively for the onset of zygotic genome activation (ZGA), and processes that dictate ZGA could also apply to H3K9me. One model postulates that cell counting mechanisms such as the amount of DNA in the embryo could be a timer (Dekens et al., 2003; Newport and Kirschner, 1982a). A second model focuses on the changes in the nuclear-to-cytoplasmic ratio as embryonic cells divide, which could be critical for diluting or concentrating a maternal repressor/activator (Newport and Kirschner, 1982b; Pritchard and Schubiger, 1996). A third model predicts that fertilization initiates a process that builds gradually as a direct readout of time passed (Ferree et al., 2016; Howe and Newport, 1996; Kimelman et al., 1987; Treen et al., 2018; Yuan et al., 2016). We tested whether any of these models could explain timing of H3K9me2 onset.

To determine if cell counting is important, we uncoupled the number of cells produced over a given unit of time. We inactivated *div-1*, a subunit of DNA polymerase-alpha primase complex, by using a temperature-sensitive mutation *(or148)* to extend the duration of S-phase and slow down cell divisions (Encalada et al., 2000). One hour after the 2-cell stage, wild-type embryos had 25–30 cells, whereas *div-1* mutant embryos had 5–15 cells (**Figure 5A**). *div-1* embryos exhibited precocious H3K9me2 based on cell number, but wild-type levels based on time post-fertilization (**Figure 5B, C**). The amount of H3K9me2 per nucleus was similar in 25–30 cell wild-type embryos and 5–15 cell *div-1* embryos. These delayed *div-1* cells had a similar volume to wild-type embryos with the equivalent number of cells (**Figure 5D**) and comparable amounts of DNA as wild-type embryos that contain the same number of cells (**Figure 5E**). This result rules out some counting models, specifically the nuclear-to-cytoplasmic ratio, the number of cells or the amount of DNA. In short, it is possible to acquire high levels of H3K9me2 without reaching a certain cell number, nuclear-to-cytoplasmic ratio or undergoing a certain number of cell divisions. Instead, this result suggests that the timing of H3K9me2 is dictated by an absolute clock that measures the time spent in interphase (∼1 hour after the 2-cell stage).

One concern was that the replicative stress or DNA damage in early *div-1* embryos caused higher levels of H3K9 methylation to engage DNA repair pathways (Ayrapetov et al., 2014). We hypothesized that if DNA damage in *div-1* mutants caused the increase in H3K9me2, one might expect restoring the faster cell cycle in *div-1* mutants to lead to even more DNA damage and H3K9me2 (**Figure 5F**). Alternatively, if the amount of time from fertilization was the critical parameter, then restoring the faster cell cycle to *div-1* mutants would rescue normal timing of H3K9me2 accumulation. To restore faster cell cycle progression to *div-1* mutant embryos, we inactivated the ATR related gene *atl-1* which leads to faster cell cycles but potentially more DNA damage (Brauchle et al., 2003). *atl-1*(RNAi); *div-1* double mutants partially suppressed the increase in H3K9me2 (**Figure 5G-I**), suggesting precocious H3K9me2 in *div-1* mutants was not due to DNA damage. Instead, timing of H3K9me2 depends on the amount of time from fertilization.

**Figure 5.**
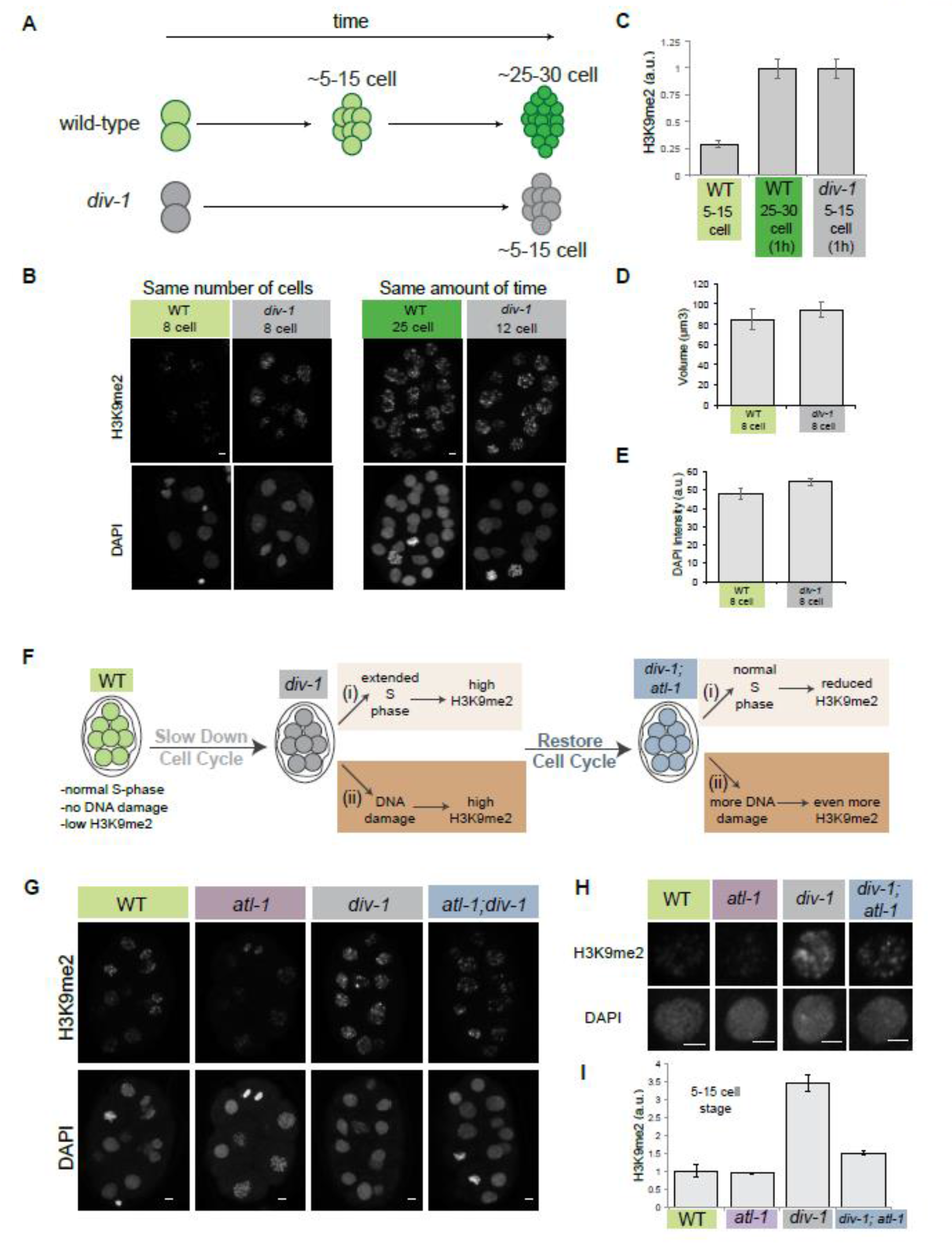
Slowing down S-phase leads to precocious accumulation of H3K9me2, based on amount of time spent after fertilization. A. Time-line of embryogenesis with respect to absolute amount of time and number of cells per embryo in wild-type (green) or *div-1(or148)* mutant (gray) embryos. B-C. H3K9me2 staining in embryos that contain the same number of cells (8-cell, left panels) or that have developed for the same amount of time after the 2-cell stage (1 hour, right panels). Scale bar 2 μm. Amount of H3K9me2 tracks with absolue time, not number of cells. D. Nuclear volume in wild-type vs. *div-1(or148)* embryos at the 8-cell stage. E. Mean DAPI intensity in wild-type vs. *div-1(or148)* embryos at the 8-cell stage. F. Rationale. Increased H3K9me2 in *div-1* mutants could be due to I) the extension of S-phase (light orange) or II) DNA damage (dark orange). Accordingly, restoring the cell cycle by inactivating *atl-1* in *div-1* mutants is predicted to have two alternative outcomes: I) rescue of cell cycle timing and reduction in H3K9me2 (light orange) or II) additional DNA damage and increased H3K9me2 (dark orange). G. H3K9me2 staining in wild-type vs. *atl-1 RNAi, div-1(or148)* and *atl-1 RNAi; div-1*(or148) double inactivation embryos at the pre-gastrula stage. Color code: wild-type – green, *atl-1* – purple, *div-1* – gray, *atl-1;div-1* – blue. In each experiment, wild-type embryos containing zen-4::gfp were included as on-slide controls (not shown). H-I. Single nuclei in interphase showing H3K9me2 levels and quantitation. Note that *atl-1* RNAi partially rescues H3K9me2 levels in *div-1* mutants in support of Scenario I (light orange from A).

Our previous work revealed that MET-2 is initially partitioned between the nucleus and cytoplasm, but becomes concentrated within nuclei as embryos mature (**Figure 6A**, (Mutlu et al., 2018)). Nuclear MET-2 is rate-limiting for H3K9me2 levels and gradually accumulates in nuclei with its binding partners LIN-65 and ARLE-14 (Mutlu et al., 2018). We wondered if precocious H3K9me2 in *div-1* mutants might reflect precocious accumulation of the MET-2 complex in nuclei. Indeed, MET-2 and its binding partners each accumulated in the nucleus earlier compared to wild-type embryos (**Figure 6B-G**). We hypothesize that the cumulative time spent in interphase after fertilization permits relocalization and activation of MET-2, and thereby dictates the onset of H3K9me2.

**Figure 6.**
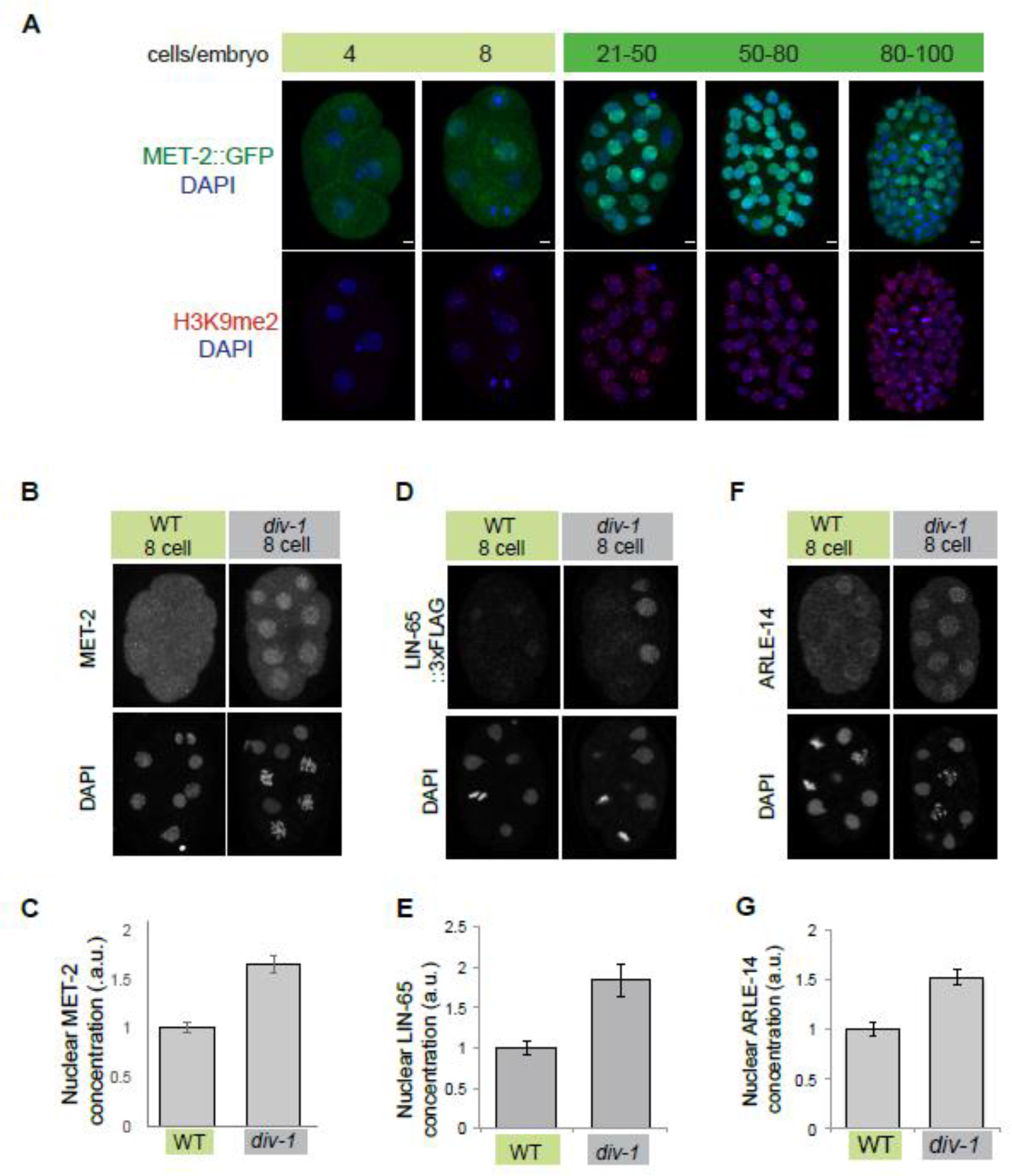
Slowing down S-phase leads to precocious accumulation of MET-2 and binding partners in the nucleus. A. MET-2::GFP localization and H3K9me2 levels during embryogenesis. MET-2::GFP (green), H3K9me2 (red), DAPI (blue), Scale bar 2 μm. Note that MET-2::GFP accumulates in the nucleus as H3K9me2 domains are formed. B,C. Wild-type vs *div-1 (or148)* embryos at the pre-gastrula stage stained with an antibody against endogenous MET-2 on the same slide. Quantitation of protein levels per nucleus. D,E. Progeny of *lin-65::3xflag* worms fed with *empty vector* vs *div-1 RNAi*. Embryos at the pre-gastrula stage were stained with a FLAG antibody. Quantitation of protein levels per nucleus. F,G. Wild-type vs *div-1 (or148)* embryos at the pre-gastrula stage stained with an antibody against endogenous ARLE-14 on the same slide. Quantitation of protein levels per nucleus.

## Discussion

This study has made three contributions to understand the timing and function of H3K9me during embryogenesis. First, the methyltransferase MET-2 promotes loss of developmental plasticity through H3K9me1/me2, and formation of higher-order heterochromatin through H3K9me3. Second, several models cannot account for the onset of MET-2 activation, including cell counting, the nuclear-to-cytoplasmic ratio or zygotic transcription. Third, the cumulative amount of time early embryonic cells spend in interphase, specifically S phase, influences accumulation of MET-2 in the nucleus and determines the onset of H3K9 di-methylation.

### Distinct roles for di vs. tri methylated H3K9 in cell fate potential and formation of higher-order heterochromatin

By sequence, MET-2 is most similar to vertebrate SETDB1(Poulin et al. 2005; Andersen and Horvitz 2007). Both enzymes can catalyze mono and di-methylation on H3K9, and directly or indirectly regulate H3K9me3 (Basavapathruni et al., 2016; Loyola et al., 2009; Towbin et al., 2012; Wang et al., 2003). SET domain methyltransferases contain a ‘switch position’ in their catalytic site that determines the degree of methylation, with bulkier residues able to accommodate mono- and di- but not tri-methylation (Jih et al., 2017). MET-2 and SETDB1 have a bulky Tryptophan residue in the switch position, which suggests that these enzymes favor mono- and di-methylation in the absence of regulatory partners. However, the presence of partially redundant H3K9 methyltransferases in mammals (including SUV39h1/2, G9a and PRDM2/3/16) and in some cases lethal phenotypes associated with loss of individual enzymes have made it hard to dissect the roles for di- vs. tri-methylation. In worms, the differential effects of the two H3K9 methyltransferases *met-2* and *set-25* on mono, di and tri methylated H3K9 provide a means to distinguish the roles of these histone marks.

Developmental potential was inversely correlated with H3K9me2 (**Figure 1, Figure 2C**): *met-2*, wild-type and *set-25* embryos had low, average and high levels of H3K9me2, respectively. *met-2* mutants were able to alter their developmental fate robustly, wild-type embryos less so, and *set-25* mutants least of all. Double mutants between *met-2* and *set-25* lacked H3K9me2, and as predicted, they extended plasticity like *met-2* single mutants. At later stages, *met-2* mutants were resistant to the CFC assay, suggesting that H3K9me2 is important for the timely loss of plasticity but not absolutely required. Additional regulators such as Polycomb and Notch may account for the partial requirement (Djabrayan et al., 2012; Joshi et al., 2010; Yuzyuk et al., 2009) We note that our results do not rule out a role for H3K9me1, but that H3K9me1 levels did not correlate with plasticity as closely as H3K9me2 levels.

EDRs were reduced in *met-2* mutants and absent in *set-25* embryos. Similarly, H3K9me3 was reduced in *met-2* and absent in *set-25* mutants, making H3K9me3 an excellent candidate to mediate higher-order heterochromatin. An intriguing but speculative idea is that emergence of EDRs depends on H3K9me3 quantitatively rather than responding to an all-or-nothing threshold (Larson et al., 2017; Strom et al., 2017).

### Higher-order heterochromatin and loss of developmental plasticity

An absence of heterochromatin could provide a permissive environment to transcribe diverse genes, an important feature of pluripotent embryos. However, we note that neither loss of EDRs in *set-25* mutants, nor absence of all H3K9me in *met-2; set-25* double mutants, did not enable HLH-1 to activate muscle genes ubiquitously. Therefore, additional mechanisms must exist to curtail the ability of cells to respond to a heterologous regulator. For example, H3K27me3 is present in *met-2*; *set-25* mutants (Towbin et al., 2012) and could compensate for the loss of H3K9 methylation. Alternatively, the local deposition of nucleosomes over transcription factor binding sites or within transcribed regions could block plasticity (Gaspar-Maia et al., 2011). In vertebrates, H3K9me3–rich domains can interfere with transcription factor binding to block reprogramming of induced pluripotent stem cells (Soufi et al., 2012). This result suggests that ‘reverse development’, as seen in reprogramming, may be qualitatively different from ‘forward development’, when plasticity is lost during embryogenesis. For example, H3K9me3 may initiate processes that eventually solidify a cell fate and block reprogramming in differentiated cells, but these processes may not yet be complete in gastrula-stage embryos.

### Zygotic transcription is not rate-limiting for H3K9me2

RNAi pathways are critical to target heterochromatin machinery in worms and other organisms such as *S. pombe* (Moazed, 2009). In particular, the piwi pathway and the nuclear RNAi (nrde) pathway both target repeat sequences that make up the bulk of heterochromatin and are marked by H3K9me2 in *C. elegans* (Ashe et al., 2012; McMurchy et al., 2017; Ni et al., 2014; Weick and Miska, 2014). While it is known that the RNAi machinery targets H3K9 methylation, it was not clear whether these components dictate the timing of H3K9me2 in embryos.

The nrde pathway acts co-transcriptionally and inhibits Polymerase II during the elongation phase of transcription through the deposition of H3K9me (Guang et al., 2010). Concomitant with a big wave in zygotic transcription, embryonic nuclei acquire methylated H3K9me (**Figure 3A**), suggesting transcription elongation could be rate-limiting for H3K9me2 deposition during embryogenesis. However, H3K9me2 levels were not affected after blocking transcription elongation (**Figure 4**), suggesting that it is not the rate-limiting component. We note that while transcription elongation was undetectable under our experimental conditions, abortive transcription may still occur. Thus, our results do not rule out Pol II transcription or RNAi as a mechanism for targeting H3K9 methylation but rule them out as timers. An intriguing model is that the absence of H3K9 methylation in early embryos provides a window of opportunity where repetitive sequences can be transcribed and be targeted by the RNAi machinery to acquire H3K9me (McMurchy et al., 2017; Penke et al., 2016; Yu et al., 2013; Zeller et al., 2016).

### Zygotic transcription and chromatin organization

The nature of the relationship between zygotic transcription and chromatin regulation has been uncertain. Chromatin regulation could precede and initiate zygotic transcription (Meier et al., 2018; Østrup et al., 2013). Conversely, zygotic transcription may help shape chromatin structure, a model that has been tested by several studies: A study that analyzed the spatial genome organization in fly embryos revealed that early expressed genes correlate with the boundaries of Topologically Associating Domains (TADs). However, emergence of these structures are not dependent on transcription (Hug et al., 2017). Another study in *Xenopus* embryos mapped the distribution of histone modifications across the genome, and found that H3K4me3 and H3K27me3 domains were established independently of zygotic transcription (Hontelez et al., 2015). Similarly, our results suggest Pol II transcription is not rate-limiting for the establishment of H3K9me in early embryos.

### Classical models of time-keeping and H3K9 methylation

Classical models of embryonic time-keeping include i) an “absolute clock” that depends on the time from fertilization (Howe and Newport, 1996; Treen et al., 2018), ii) the increasing nuclear-to-cytoplasmic ratio (Newport and Kirschner, 1982b; Pritchard and Schubiger, 1996) or iii) cell counting via increasing DNA content in the embryo, which titrates a maternal repressor (Dekens et al., 2003; Newport and Kirschner, 1982a). These models have mostly been studied in the context of zygotic genome activation. One study analyzed local changes in chromatin organization during fly embryogenesis and found that promoter accessibility is controlled by the nuclear-to-cytoplasmic ratio (Blythe and Wieschaus, 2016). However, large-scale changes in chromatin organization during development such as heterochromatin formation had not been studied through the lens of these models.

In this study, we applied these models to H3K9 methylation, which emerges during gastrulation (Mutlu et al., 2018). Our analysis of *div-1* mutants ruled out cell counting as a potential timer and suggests that H3K9me2 depends on an absolute clock for nuclear entry of MET-2 and its binding partners. *div-1* mutants specifically extend S-phase compared to wild-type embryos (Encalada et al., 2000), suggesting that the cumulative time that early embryonic cells spend in S-phase is critical for measuring time.

### Spatial regulation of MET-2 and timing of H3K9 methylation

MET-2 moves gradually into nuclei in the pre-gastrula embryo, but is released into the cytosol during mitosis (Mutlu et al., 2018). Early embryonic cells divide rapidly, with a 40-minute cell cycle that likely interferes with the accumulation of MET-2 in nuclei. Slowing down the cell cycle in *div-1* embryos leads to precocious accumulation of MET-2 in the nucleus (**Figure 6**). Consistent with this idea, cells that divide slowly compared to other cell lineages (e.g. the E cells (Sulston et al., 1983)) accumulate more nuclear MET-2 and more H3K9me2 than rapidly dividing cells (data not shown). In sum, our data suggest that the cumulative time early embryonic cells spend in interphase could be important for i) MET-2 accumulation in the nucleus and ii) deposition of H3K9me by nuclear MET-2.

Spatial regulation of proteins provides a rapid means to restrict their activity and is a common theme in temporal regulation of developmental processes. For instance, Prmt1 and SIRT1 each transition from the cytosol to the nucleus, or vice versa, to alter their activity upon differentiation (Ancelin et al., 2006; Hisahara et al., 2008). Similarly, the onset of zygotic transcription is dictated by the OMA proteins, which sequester a critical cofactor for TFIID in the cytoplasm (Guven-Ozkan et al., 2008). Our study provides a third example, where the embryonic clock for H3K9me depends on the spatial regulation of MET-2 and its binding partners.

## Materials and Methods

### Strains

Strains were maintained at 20°C according to (Brenner, 1974), unless stated otherwise*.

*N2* (wild-type Bristol)
*EU548 *div-1(or148ts)* III. (Encalada et al., 2000)
*KW1975 *taf-6.2(ax514); unc-17(e113)* IV.
SM2440 *jmjd-1.2 (tm3713)* IV. (Kleine-Kohlbrecher et al., 2010)
EL597 *omIs 1 [Cb-unc-119 (+) met-2::gfp* II].
SM2575 *lin-65::3xflag* I.
SM2333 *pxSi01 (zen-4::gfp, unc-119+)* II*; unc-119(ed3)* III.
KM167 *HS::hlh-1*, (Fukushige and Krause, 2005).
SM1623 *HS::hlh-1; met-2 (ok2307*) III.
JAC500 *his-72(csb43[his-72::mCherry])* III. (Norris et al., 2015)

### Antibody staining

Antibody staining was performed as described previously (Mutlu et al., 2018). The following antibodies were used for immunostaining by 5min 2% paraformaldehyde (PFA), 3 min methanol (for all except PHA-4, which was fixed with 10min 2% PFA, 3min methanol). H5 Pol II staining followed a completely different protocol described in (Kaltenbach et al., 2000).

H3K9me2 (1:200) Abcam ab1220, MABI0307 Kimura 6D11
Histone H3 (1:500) Abcam ab1791
Pan-histone (1:500) Chemicon/Millipore MAB052
FLAG M2 (1:100) Sigma F1804
MET-2 (1:500) Raised against the first 17 amino acids of MET-2 and affinity purified, a gift from Eleanor Maine.
ARLE-14 (1:500) Generated by our lab as described in (Mutlu & Mango, submitted).
H5 Polymerase II (1:100) Covance MMS129-R
Paramyosin (1:50) Developmental Studies Hybridoma Bank 5–23
PHA-4 N-terminus (1:1000) (Kaltenbach et al., 2005)

### TEM

was done as described previously (Mutlu et al., 2018).

### Quantitation of histone modifications and nuclear proteins

Analysis was done as described in (Mutlu et al., 2018). Briefly, embryos were imaged with a ZEISS LSM700 or LSM880 Confocal Microscope and analyzed by Volocity Software. Signal intensity of marks were calculated for each nucleus and average values for nuclei at designated embryonic stages plotted.

### Transcription blocking

KW1975 *taf-6.2(ax514)* was maintained at 15°C. For experiments, KW1975 L4s were fed bacteria containing an *ama-1 RNAi* vector. In parallel, SM2233 L4s were fed with bacteria containing *empty vector*. Both strains were grown at 15°C for 50–60 hours. Adult worms were dissected at 26°C and 1–4 cell embryos were transferred onto the same poly-L-lysine slide. Embryos were aged for 1 hour at 26°C in a humidity chamber and stained for H3K9me2. For H5 Polymerase II stains, wild-type JAC500 worms fed with *empty vector* were used as an on-slide control instead of SM2233.

### Pouring RNAi plates

RNAi clones from the Ahringer library were used unless stated otherwise. First, identity of clones was confirmed by sequencing. To pour plates, bacteria were grown in 5ml LB with 5μl Carbenicillin (100mg/ml) for 6–8 hours at 37°C and pelleted at 4000rpm for 10 minutes. The bacterial pellet was resuspended in 400μl 0.5M IPTG, 30μl 100mg/ml Carb and 70μl Nuclease-free water. 5ml NGS plates were seeded with 100μl of resuspended bacterial solution and kept at room temperature for 2 days before use.

### *div-1* experiments

*div-1(or148ts)* was maintained at 15°C and shifted to 26°C for experiments. 2-cell stage wild-type *zen-4::gfp* (SM2233) and *div-1* (EU548) embryos were picked and aged for 1 hour on the same poly-L-lysine slide in a humidity chamber at 26°C. H3K9me2 staining after embryonic temperature shifts compared 25–30 cell wild-type embryos to 10–15 cell *div-1* embryos.

Temperature shifts to 26°C that started with L4 animals instead of embryos produced identical results in terms of H3K9me2 levels at given embryonic stages. In L4-shift experiments, mixed stage SM2233 and EU548 embryos were dissected from gravid adults and stained for H3K9me2 on the same slide. Cells that contained the same number of cells were compared using L4-shifts.

For *atl-1* rescue experiments, *N2* L4s were fed with *atl-1 RNAi* bacteria. *div-1(or148ts)* L4s were fed with either *empty vector* or *atl-1 RNAi* bacteria overnight at 26°C. SM2233 L4s were fed with bacteria containing an *empty vector* at 26°C and included as an on-slide control on all experiments. Embryos were dissected from gravid adults and stained for H3K9me2.

### CFCA

Cell Fate Challenge Assay was conducted similarly to (Kiefer et. al, 2007). In brief, two-cell embryos were collected from wild-type, *set-25* or *met-2* mothers carrying an integrated *HS::hlh-1* array (Fukushige and Krause, 2005). Embryos were incubated at 20°C for 3 hours until they reached the 100-cell stage, determined by DAPI staining and cell counts. Heat shock was administered at 33°C for 30 minutes on Poly-L-Lysine slides in a humidity chamber and embryos were incubated at 20°C for 20 hours. Terminally differentiated embryos were stained for paramyosin (muscle, (Fukushige and Krause, 2005)) and PHA-4 (foregut, (Horner et al., 1998)). Embryos were imaged using the Zeiss LSM 700 confocal microscope. RNA expression analysis for hlh-1 was done as described previously (Mutlu et al., 2018).

### GO Term Analysis

H3K9me1 (GSE49744), H3K9me2 (GSE49736) and H3K9me3 (GSE49732) methylated regions were defined by a MACS2 broad peak call (Zhang et al., 2008) and the center of peaks were assigned to genes to curate a list. DAVID (https://david.ncifcrf.gov/) was used for GO Term analysis.

## Acknowledgements

We thank the Harvard Center for Biological Imaging, K. Nguyen and B. Raja for their help on TEM procedures. We acknowledge support from the NIH (R37GM056264), the John D. and Catherine T. MacArthur Foundation and Harvard University to SEM, the AAUW International Fellowship to BM. Some strains were provided by the Caenorhabditis Genetics Center, funded by NIH P40OD010440. All data needed to evaluate conclusions in the paper are present in the paper and the Supplementary Materials. Additional data available from authors upon request. Reagents can be provided by Susan Mango pending scientific review and a completed material transfer agreement.

## Author contributions

BM and SEM designed and conducted the study and wrote the manuscript. DHH performed TEM. HMC performed GO Term analysis for H3K9me. Authors have no competing interests.

